# BrainYears: A functional EEG-based brain age clock enables intervention-ready measurements of brain aging

**DOI:** 10.64898/2026.03.26.714124

**Authors:** Sierra Lore, Corey Julihn, Paola Telfer, Morten Scheibye-Knudsen, Eric Verdin

**Affiliations:** Buck Institute for Research on Aging, 8001 Redwood Blvd, Novato, 94945, California, USA; University of Copenhagen, Nørregade 10, 1172 København, Denmark; 1055 Dunsmuir Street, Suite 3000, Vancouver, BC, Canada V7X 1K8

**Keywords:** brain aging, brain age clock, biological age, electroencephalography (EEG), event-related potential (ERP), neuromodulation, brain biomarkers, neurotechnology

## Abstract

Biological brain aging is a major determinant of cognitive decline and neurodegenerative disease, yet scalable and intervention-ready brain aging biomarkers remain limited. Here, we develop an electroencephalography (EEG)-based brain age clock using machine learning trained on high-dimensional neural features across the adult lifespan. Using 643 features captured from a Sens.ai headset and controller device, the model predicts chronological age with high accuracy (Pearson r = 0.92; MAE = 4.43 years) and yields an interpretable set of age-informative neural features capturing functional signatures of brain aging. Unlike MRI-based approaches, this EEG-based clock is non-invasive, transportable, cost-effective and suitable for repeated at-home longitudinal measurement. Furthermore, in a longitudinal neuromodulation program, BrainYears-predicted brain age decreased by a mean of −5.18 years in the intervention group whereas a minimal-exposure comparison group showed no change on average (+0.07 years). Together, this work introduces a functional brain aging biomarker and an intervention-ready platform for quantifying brain age modulation.

---

Aging of the human brain is accompanied by progressive changes in neural function that contribute to cognitive decline, increased disease susceptibility and reduced capacity for plasticity^1–3^. These changes are highly heterogeneous across individuals, such that chronological age alone is a poor predictor of brain health, cognitive performance or vulnerability to neurodegenerative disease^4,5^. This heterogeneity has motivated the development of biomarkers that aim to quantify biological rather than chronological aging^6,7^. Brain age clocks, which estimate chronological age from neurobiological data using statistical or machine-learning approaches, have emerged as a particularly informative class of biological age biomarkers^8,9^. The deviation between predicted brain age and chronological age—often referred to as “brain age acceleration”—has been associated with Alzheimer’s disease^10^, Parkinson’s disease^11^, psychiatric disorders^12^, traumatic brain injury^13^ and increased mortality risk^14^. These findings support the biological relevance of brain age as a measure of neural aging beyond time alone^15,16^.

To date, the majority of brain age clocks have been derived from structural magnetic resonance imaging (MRI), using features such as cortical thickness, gray-matter volume and white-matter integrity^15,17^. MRI-based clocks have demonstrated strong predictive performance and clear clinical associations. However, their reliance on expensive, infrastructure-intensive imaging limits scalability, accessibility and feasibility for frequent longitudinal assessment^18–20^. These constraints pose a major barrier for applications that require dense sampling, population- scale deployment, or repeated measurements to evaluate interventions.

Electroencephalography (EEG) provides a complementary and underutilized substrate for quantifying brain aging. EEG directly measures neural activity at millisecond temporal resolution and captures functional properties of brain networks, including oscillatory dynamics, synchronization, and temporal variability^21–23^. Event-related potentials (ERPs), computed by averaging time-locked responses to task stimuli, isolate transient components such as the N100, P200, N200, and P3, which index sensory processing, attentional allocation, conflict monitoring, and response inhibition during cognitive control tasks. These properties are shaped by synaptic integrity, excitation–inhibition balance and large-scale network coordination—processes that are known to change with aging and that may plausibly respond to intervention^24^. Moreover, EEG is non-invasive, comparatively low-cost and readily deployable outside specialized imaging centers, making it well suited for large-scale and longitudinal studies^25^.

An additional limitation of most existing brain age clocks, regardless of modality, is their predominantly observational framing. While accelerated brain aging has been robustly associated with disease and adverse outcomes, it remains unclear whether brain age represents a fixed trajectory or a modifiable biological state^6,26^. Addressing this question requires biomarkers that are not only predictive but also suitable for repeated measurement and capable of detecting subtle functional changes in response to intervention. Such biomarkers would enable direct testing of whether targeted interventions can slow, halt or reverse aspects of brain aging.

Here, we develop BrainYears, a scalable EEG-based brain age clock explicitly designed to address these gaps^25,27^. Using machine learning trained on high-dimensional EEG-derived features and demographic covariates across the adult lifespan, BrainYears predicts chronological age with high accuracy. The architecture and deployment context of this Sens.ai device and clock are optimized for longitudinal and interventional use. EEG-based measurement enables repeated sampling within individuals over short time scales, facilitating the study of brain age dynamics rather than static estimates. Together, this work introduces an accessible and interpretable biomarker of functional brain aging and provides an intervention-based platform for probing the plasticity of brain aging in translational and therapeutic contexts.

## Results

### A deployable EEG platform for longitudinal brain-age quantification

We developed BrainYears, a scalable EEG-based brain-age clock designed for repeated, real-world deployment using the Sens.ai neurotechnology platform (Fig. 1a). The integrated system comprises a wearable EEG headset, a GeniusPulse controller, and a synchronized mobile application enabling standardized assessment and intervention workflows. The headset incorporates midline EEG sensors positioned at Fz, Cz, and Pz (Fig. 1b), seven 810-nm near-infrared (NIR) LEDs for transcranial photobiomodulation (Fig. 1c), and a pulse oximeter positioned near the left ear (Fig. 1d), enabling simultaneous electrophysiological acquisition and physiological modulation.

**Figure 1.**
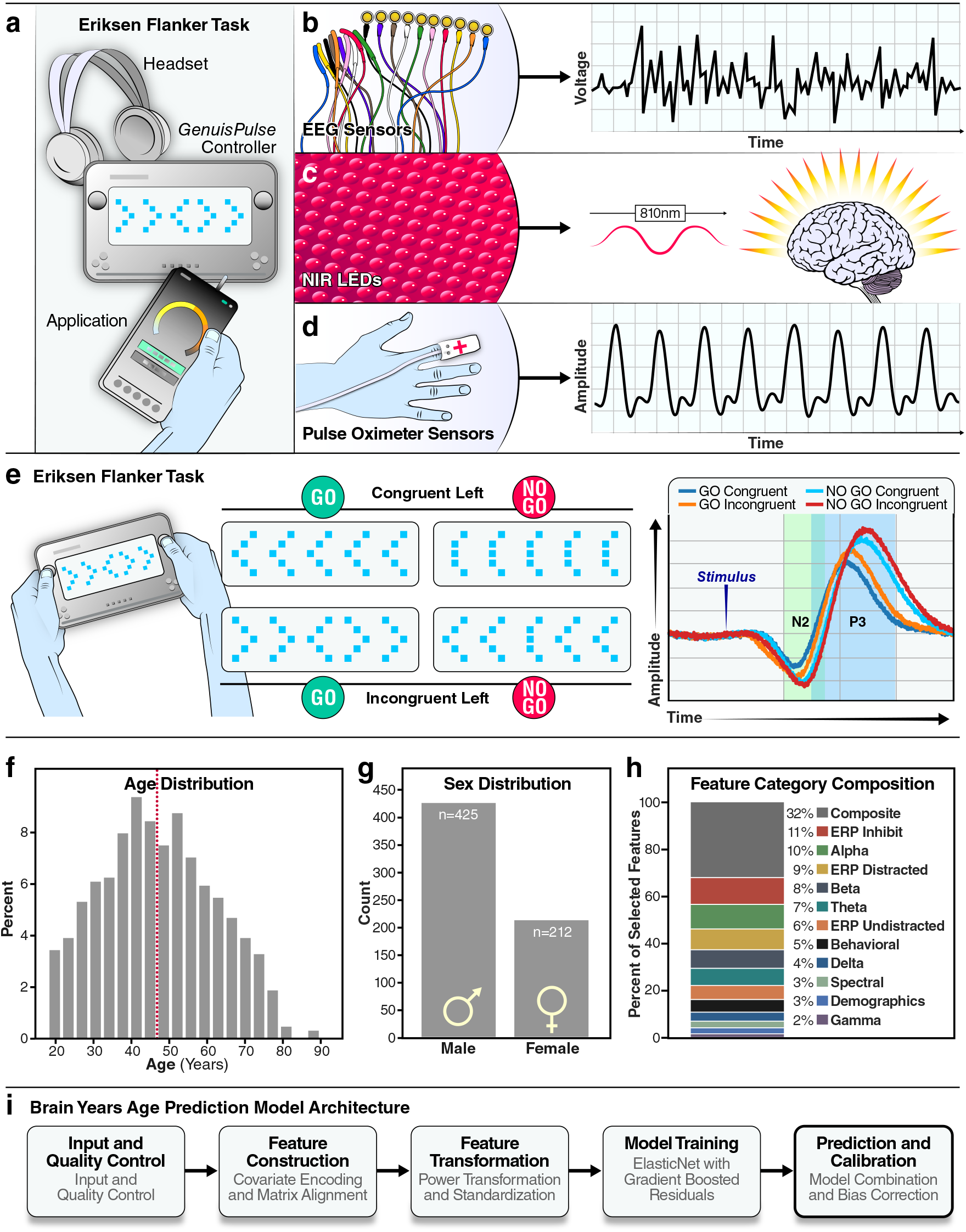
Platform, cohort and modeling framework for EEG-based brain age prediction. **a**, Novel device system including wearable EEG headset, GeniusPulse controller and companion mobile application. **b**, Midline EEG electrodes (Fz, Cz, Pz) used during ERP acquisition; representative voltage trace shown. **c**, Displaying the 7 near-infrared LEDs at the 810 nm wavelength used during neuromodulation intervention sessions. **d**, Peripheral pulse oximeter sensor and representative amplitude trace. **e**, Eriksen flanker task with embedded no-go trials; canonical N2 and P3 global averaged wave forms are shown across trial types. **f**, Chronological age distribution of participants included in model development; data were split 80:20 into training and held-out test sets prior to preprocessing. **g**, Sex distribution; sex was one-hot encoded using training-set parameters. **h**, Proportion of features by category within the 643- feature matrix used for modeling. Categories include ERP Inhibit (correct inhibit and error- related ERP biomarkers), ERP Undistracted (correct congruent trials), ERP Distracted (correct incongruent trials), canonical EEG bands (delta, theta, alpha, beta, gamma), Behavioral (reaction time and accuracy measures), Composite (engineered interaction features), Demographics (race, sex, education), and additional Spectral features (e.g., broadband slope, total power, median frequency). **i**, BrainYears modeling framework: variance filtering and percentile-based clipping, Yeo–Johnson transformation and standardization, ElasticNet regression with gradient-boosted residuals, and polynomial age-bias correction; all preprocessing was fit on training data only.

Functional brain assessment was performed using a standardized Eriksen flanker task with embedded go/no-go trials (Fig. 1e), capturing stimulus processing, cognitive control, conflict monitoring, and inhibitory function. ERP segments (Go congruent, Go incongruent, No-go congruent, No-go incongruent) were artifact filtered and averaged prior to feature extraction. The modeling cohort spanned early adulthood to late life (Fig. 1f), included both sexes (Fig. 1g), and reflected diversity in race and educational background (Supplementary Fig. S1a,b).

From these recordings, we used 643 features outputted from the Sens.ai device. For interpretability, features were grouped into predefined physiological and task-based categories. ERP Inhibit refers to ERP biomarkers derived from correct inhibit trials and error-related ERP biomarkers; ERP Undistracted refers to ERP biomarkers derived from correct undistracted trials; and ERP Distracted refers to ERP biomarkers derived from correct distracted trials. Canonical EEG band categories (delta, theta, alpha, beta, gamma) correspond to band-annotated oscillatory biomarkers. Behavioral features comprise reaction time and accuracy-based measures. Composite features represent engineered interaction measures derived from combinations of primary signals. Demographic features include race, sex, and education covariates. Spectral features refer to additional spectral summary measures such as total power, broadband slope, or median frequency (Fig. 1h).

### BrainYears modeling captures linear and nonlinear structure in functional aging

BrainYears implements a two-stage machine-learning architecture designed to disentangle dominant linear aging effects from nonlinear higher-order structure. ElasticNet regression first learned sparse, regularized linear associations between features and chronological age. A gradient-boosted regressor was trained on the residuals using percentile-clipped features and the ElasticNet prediction as inputs, capturing nonlinear structure beyond the linear age signal. All preprocessing—including variance filtering, percentile clipping, Yeo–Johnson transformation, and standard scaling—was fit exclusively on training data and serialized within a deployable model bundle to ensure deterministic inference across longitudinal applications.

On a held-out test set (n = 128; 20%), predicted age closely tracked chronological age (Pearson r = 0.92; R^2^ = 0.85; MAE = 4.42 years; Fig. 2a). The first principal component of the feature space explained 28.6% of total variance and correlated strongly with chronological age (r = 0.68; Fig. 2b), indicating that age-related variance occupies a dominant, coherent axis within the electrophysiological manifold. Calibration analysis after polynomial bias correction demonstrated near-linear agreement across the adult lifespan (calibration slope = 1.01, intercept = −0.28; Fig. 2c). Mean absolute error remained stable across age bins (4.83 ± 0.95 years; Fig. 2d), and residuals were centered near zero (mean = −0.29 years) with minimal systematic drift (Fig. 2e). Together, these results confirm that chronological age is robustly encoded in distributed electrophysiological features spanning oscillatory, evoked, and network domains.

**Figure 2.**
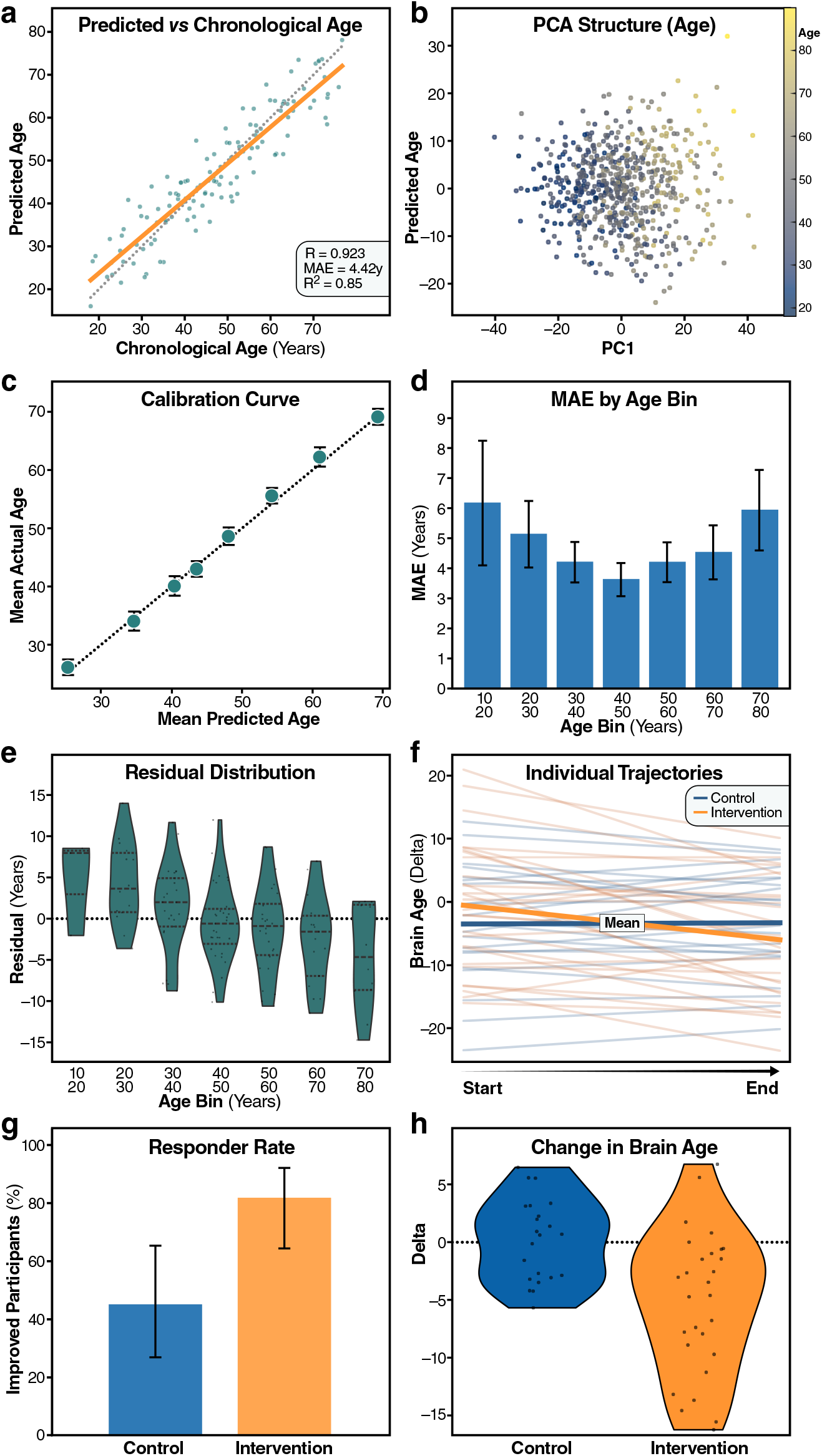
BrainYears prediction performance and longitudinal modulation. **a**, Predicted versus chronological age in the held-out test set after bias correction (Pearson r = 0.92; R^2^ = 0.85; MAE = 4.42 years). **b**, Principal component analysis of the feature space colored by age. **c**, Calibration curve showing mean predicted versus mean actual age across bins after bias correction. **d**, Mean absolute error stratified by age bin. **e**, Distribution of prediction residuals across age bins. **f**, Individual longitudinal brain-age trajectories (pre to post); thin lines represent participants and bold lines represent group means. **g**, Percentage of participants demonstrating improvement (negative brain-age delta) in intervention and minimal-exposure comparison groups. **h**, Distribution of pre–post brain-age delta by group (intervention mean −5.18 years; comparison mean +0.07 years).

Importantly, the BrainYears model bundle was serialized and reused without retraining for all downstream inference, enabling strict separation between model development and unbiased longitudinal evaluation.

### Brain age is dynamically measurable within individuals

To evaluate longitudinal sensitivity, we applied the frozen BrainYears model to paired pre/post assessments surrounding a structured multi-week neuromodulation program delivered through the Sens.ai platform. Individual trajectories revealed consistent downward shifts in bias-corrected brain age following intervention (Fig. 2f). Across adherent participants, 85% exhibited improvement (Fig. 2g), with a mean reduction of −5.18 years in predicted brain age (Fig. 2h).

In contrast, a minimal-exposure comparison group demonstrated near-zero mean change (+0.07 years), providing an empirical reference for test–retest stability under low engagement conditions. The magnitude and directionality of change observed in the intervention cohort exceeded residual variability observed in the comparison group (control 95th percentile of absolute change = 5.67 years), supporting the capacity of the clock to detect within-individual functional shifts over multi-week intervals.

These findings suggest that electrophysiological brain age, as assessed by BrainYears, is not fixed but dynamically responsive over time. However, the observed reductions do not necessarily imply reversal of structural aging, but rather modulation of a functional age estimate derived from network-level electrophysiological dynamics. Given that EEG captures oscillatory coordination, excitation–inhibition balance, and task-evoked processing properties, features known to reflect synaptic and network integrity, the results are consistent with the possibility that aspects of functional brain aging may be partially modifiable.

Importantly, the magnitude of change occurred over weeks rather than years, indicating that the model is sensitive to short-interval alterations in neural function. Whether these shifts reflect durable reorganization, transient state-dependent changes, or a combination thereof will require longer-term follow-up. Nonetheless, the ability to detect coherent directional movement within individuals establishes that this EEG-derived age metric behaves as a dynamic functional biomarker rather than a static chronological correlate.

### Functional brain aging reflects coordinated domain-level reorganization

We next examined the structure of age-associated signal within BrainYears across feature domains. Category-level distributions of feature–age correlations demonstrated that aging information is broadly distributed across composite, ERP, oscillatory, behavioral, spectral, and demographic domains rather than concentrated in a single marker class (median absolute correlation range across categories = 0.04–0.51; Fig. 3a). Permutation importance analysis further confirmed this distributed architecture, with composite features exerting the largest contribution (ΔMAE = 5.70 years), followed by ERP Inhibit (1.21 years), ERP Undistracted (0.90 years), ERP Distracted (0.80 years), alpha-band features (0.73 years), and behavioral metrics (0.72 years) upon category disruption (Fig. 3b). Notably, all evaluated domains produced positive ΔMAE values, indicating non-redundant contributions to predictive performance.

**Figure 3.**
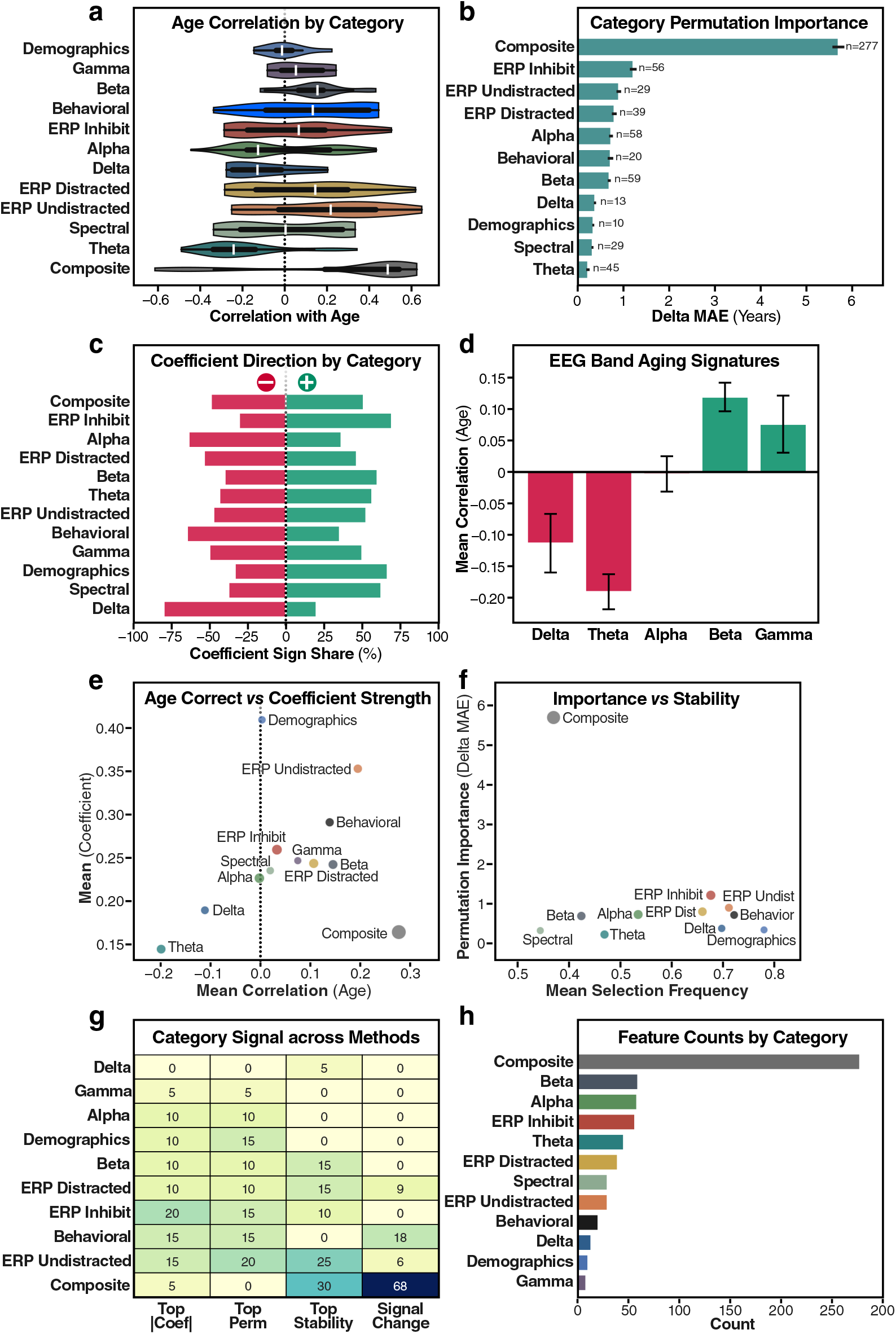
BrainYears domain-level feature structure and aging signatures. **a**, Distribution of univariate feature correlations with chronological age stratified by functional category, showing that age-associated signal is distributed across composite, ERP, oscillatory, behavioral, spectral and demographic domains. **b**, Category-level permutation importance expressed as the increase in mean absolute error (ΔMAE, years) upon shuffling features within each domain; larger ΔMAE indicates greater predictive contribution. **c**, Share of positive and negative ElasticNet coefficients by category, reflecting domain-specific directionality of age associations. **d**, Mean age correlation for canonical EEG frequency bands (delta, theta, alpha, beta, gamma), highlighting reductions in slow-frequency power and increases in higher-frequency activity with age. **e**, Mean absolute coefficient magnitude versus mean age correlation by category, illustrating concordance between univariate age association strength and multivariate model weighting. **f**, Permutation importance plotted against mean feature selection frequency across resampled splits, demonstrating stability of high-impact domains. **g**, Heatmap summarizing the proportion of features from each category represented among top-ranked features by coefficient magnitude, permutation importance, selection stability and significant age change. **h**, Total number of features per category included in the modeling framework.

Coefficient directionality analysis revealed systematic patterns across domains, with distinct proportions of positive and negative age associations depending on feature class (Fig. 3c). At the frequency-band level, mean age correlations demonstrated canonical aging signatures, including negative associations in delta (mean r = −0.11) and theta (mean r = −0.19) bands and positive associations in beta (mean r = 0.12) and gamma (mean r = 0.08) bands (Fig. 3d), consistent with known shifts in oscillatory balance during aging^28^.

To assess concordance between association strength and model weighting, we compared mean feature–age correlation with mean coefficient magnitude across domains (Fig. 3e). Domains with stronger age correlations generally exhibited larger model weights, supporting biological interpretability of learned parameters. Similarly, permutation importance scaled with feature selection frequency across resampled splits (Fig. 3f), indicating that highly influential domains were also consistently retained during model fitting.

A summary heatmap integrating coefficient magnitude, permutation importance, selection stability, and significant age-change metrics further demonstrated coordinated domain-level contributions (Fig. 3g). Composite features constituted 43% of the total feature space (277 of 643 features), but ERP and oscillatory domains collectively accounted for substantial predictive structure. Median absolute pre–post effect sizes varied by domain (composite |d| = 0.43; behavioral = 0.41; Supplementary Fig. S1c), indicating heterogeneous yet coordinated remodeling of electrophysiological architecture with age. Selected feature strength distributions likewise differed across categories (Supplementary Fig. S1d), with median absolute coefficients ranging from 0.18 (beta band) to 0.44 (demographic covariates), and ERP domains exhibiting intermediate magnitudes (ERP Undistracted = 0.30; ERP Inhibit = 0.25).

Together, these analyses demonstrate that BrainYears does not rely on a single electrophysiological marker but instead integrates distributed oscillatory, task-evoked, behavioral, and composite interaction features into a coherent systems-level representation of functional brain aging.

### A scalable framework for interventional brain-aging research

Unlike imaging-based clocks constrained by sparse sampling and high acquisition cost, this EEG-based framework enables dense longitudinal measurement in ecologically valid settings. The separation of linear and nonlinear components enhances sensitivity to subtle functional deviations while preserving interpretability. The fully serialized preprocessing and bias-correction pipeline ensures reproducibility across cohorts and time points without recalibration.

Collectively, these results establish a deployable electrophysiological clock that (i) generalizes across the adult lifespan, (ii) exhibits stable calibration properties, (iii) captures distributed, domain-level aging signatures, and (iv) detects within-individual functional shifts following structured neuromodulation exposure.

This framework positions EEG-based brain-age modeling as a scalable translational tool for mechanistic aging research and intervention monitoring, bridging laboratory-grade electrophysiology with translational longitudinal deployment.

## Discussion

In this study, we introduce BrainYears, a deployable EEG-based brain-age clock that captures distributed, task-evoked and oscillatory signatures of functional aging and demonstrates measurable within-individual shifts over multi-week intervals. By integrating standardized ERP assessment with a two-stage machine-learning architecture and deterministic preprocessing, we establish a framework that bridges laboratory-grade electrophysiology with scalable, longitudinal deployment. The resulting model generalizes across adulthood, exhibits coherent structure in feature space, and encodes age information across multiple physiological domains, positioning EEG-derived brain age as a translational biomarker candidate.

A central finding of this work is that functional brain aging cannot be reduced to a single electrophysiological marker. Rather, age-associated signal is distributed across ERP-derived features from inhibit, undistracted, and distracted trials; canonical oscillatory band power (delta, theta, alpha, beta, gamma); additional spectral summary measures; behavioral performance metrics; engineered composite interaction features; and demographic covariates (Fig. 3a). Band-level signatures (Fig. 3d) and coefficient directionality patterns (Fig. 3c) indicate coordinated shifts across canonical frequency ranges and task-evoked responses, consistent with the view that aging reflects systems-level reorganization of neural timing, excitation–inhibition balance, and network integration. Highly ranked features were stable across resampled splits (Fig. 3f), suggesting that the aging signal contains a reproducible electrophysiological core rather than being driven by dataset-specific fluctuations. Concordance across modeling further supports robustness of category-level contributions (Fig. 3e).

The two-stage architecture was explicitly designed to disentangle dominant linear aging trends from nonlinear residual structure. The ElasticNet layer captures sparse linear associations, while gradient-boosted residual modeling isolates higher-order interactions not explained by linear effects. This design yields both interpretability and sensitivity. Coefficient magnitude scales with age association strength even after bias correction (Fig. 3e), indicating that learned weights reflect meaningful biological signal rather than regression artifacts. From a methodological standpoint, this separation is critical. Many age-prediction models implicitly conflate linear trend capture with nonlinear feature interaction. By modeling residuals explicitly, we capture nonlinear components of age-associated variance not explained by linear effects. This architecture is particularly important when evaluating intervention-induced shifts, which could manifest as subtle nonlinear perturbations rather than large-scale linear reversals.

The clock demonstrated strong generalization on held-out data (Fig. 2a), with stable error across age bins (Fig. 2d) and residual were centered near zero across most age bins (Fig. 2e). Principal component structure aligned strongly with chronological age (Fig. 2b), indicating that age-associated variance occupies a dominant axis within the EEG feature manifold. Post hoc polynomial bias correction yielded well-calibrated predictions across the lifespan (Fig. 2c), minimizing regression-to-the-mean effects that commonly confound brain-age models. All preprocessing parameters were fit exclusively on training data and serialized within a model bundle reused unchanged for longitudinal inference, eliminating train–test leakage and ensuring reproducibility across cohorts and time points. In contrast to models requiring retraining or recalibration for new datasets, this framework supports direct longitudinal application without drift.

A defining motivation for this work was to determine whether electrophysiological brain age can be measured dynamically within individuals. In the intervention cohort, BrainYears predicted bias-corrected age decreased by an average of 5.18 years, with 85% of participants exhibiting improvement (Fig. 2f–h). In contrast, a minimal-exposure comparison group demonstrated near-zero mean change, providing an empirical reference for test–retest stability. These findings suggest that the clock is sensitive to functional shifts occurring over multi-week intervals. Individual trajectories (Fig. 2f) demonstrate coherent directional movement rather than random fluctuation, supporting the biological plausibility of measurable functional modulation. The ability to detect within-individual shifts distinguishes EEG-based clocks from many imaging-based models optimized primarily for cross-sectional discrimination and less suited for dense longitudinal sampling. Because EEG acquisition is low-cost, portable, and repeatable, this framework enables high-frequency monitoring of functional trajectories.

The framework carries several translational implications. It establishes that distributed electrophysiological features encode sufficient age-related information to construct a robust clock without reliance on structural imaging. It further demonstrates that such a clock can be embedded within an integrated assessment–intervention platform (Fig. 1a–d), enabling repeated measurement in real-world contexts. The model’s interpretable feature space allows domain-level hypothesis generation regarding oscillatory balance, task-evoked dynamics, and network coordination. Because interventions targeting brain aging are expected to produce gradual, systems-level changes rather than abrupt structural shifts, a modality capable of capturing distributed functional remodeling is particularly well suited to interventional evaluation.

Several limitations warrant consideration. The intervention cohort was not randomized, and potential confounding factors—including sleep, medication changes, lifestyle variation, or placebo effects—were not controlled. Future studies incorporating randomized controlled designs, larger cohorts, and extended follow-up will be necessary to determine durability and specificity of observed shifts. Additionally, while distributed electrophysiological changes are detectable, the biological substrates underlying these shifts remain to be defined. Integration with structural imaging, molecular aging markers, or cognitive endpoints will be important for establishing convergent validity.

Despite these limitations, the present work supports a conceptual shift: functional brain age, measured through distributed electrophysiological dynamics, is quantifiable, reproducible, and longitudinally sensitive. By combining standardized ERP assessment, deterministic preprocessing, residual-based modeling, and serialized deployment, we define a scalable framework for monitoring functional brain aging. As efforts to modulate aging biology accelerate, biomarkers capable of detecting short-interval functional change will become increasingly important. This EEG-based clock provides a deployable candidate for such monitoring, bridging mechanistic neuroscience and translational geroscience, and enabling function-centered evaluation of brain aging trajectories.

## Online Methods

### Study design and datasets

This study comprises (i) development of an EEG-based brain age model from cross-sectional ERP assessments collected on the Sens.ai platform, and (ii) application of the trained model to paired pre/post assessments surrounding a structured at-home neuromodulation program and a minimal-exposure comparison group. Model development and evaluation were performed on a participant-level 80:20 split into training and held-out test sets, with all preprocessing fit on training data only and then applied unchanged to the held-out test data (Fig. 2a). Longitudinal analyses were performed separately using the frozen preprocessing + model bundle trained on the cross-sectional dataset.

### Ethics, consent and privacy

All analyses were conducted as a secondary analysis of existing Sens.ai customer data. Participants provided informed, voluntary consent that included use of de-identified data for scientific and/or medical research. Prior to analysis, datasets were stripped of direct identifiers and handled under secure access controls. No individual-level data are reported; results are presented only in aggregate. The dataset includes age, sex, education, and race as self-reported demographics.

### Sens.ai device system and sensors

Recordings were obtained using the Sens.ai neurotechnology platform, which integrates a wearable EEG headset, a handheld controller, and a companion mobile application (Fig. 1a). EEG was acquired from three midline scalp electrodes positioned at Fz, Cz, and Pz (Fig. 1b). The headset includes a peripheral pulse oximeter sensor positioned near the left ear (Fig. 1d). For neuromodulation sessions, the device incorporates seven near-infrared LEDs (wavelength 810 nm; power density 312 mW/cm^2^) used for transcranial photobiomodulation (Fig. 1c).

### ERP assessment task

Functional brain assessment was performed using an Eriksen flanker task with embedded go/no-go trials (Fig. 1e). Participants indicated the direction of a central arrow while ignoring flanking distractor arrows. Congruent trials presented distractors pointing in the same direction as the target; incongruent trials presented distractors pointing in the opposite direction. No-go trials required response inhibition and used a rounded central target to cue withholding a response. The protocol comprised 400 trials in total: 65 left congruent, 65 right congruent, 65 left incongruent, 65 right incongruent, 35 no-go congruent right, 35 no-go congruent left, 35 no-go incongruent right, and 35 no-go incongruent left.

### Signal processing and ERP feature extraction

Sensor recordings were processed with trial-level artifact handling, and artifact-free trials were averaged within task segments. Segments included Go Congruent, Go Incongruent, No-go Congruent, No-go Incongruent, and Commission Error trials. Canonical ERP component features (including N100, P200, N200, and P300) were extracted per segment as amplitude- and latency-based summary measures. Behavioral performance features were extracted from the task, including accuracy and reaction time (RT) metrics such as distracted accuracy and RT from incongruent trials, RT variability, inhibition accuracy, and post-error RT. In addition to ERP and behavioral features, the Sens.ai device outputs include oscillatory and spectral descriptors derived from the EEG signal (e.g., band-annotated power features and broader spectral summaries), and engineered composite interaction features computed from pairs or combinations of primary measures. The modeling described below uses the exported numeric feature matrix as provided by Sens.ai.

### Feature matrix construction and category mapping

The brain age model was trained using a fixed exported feature matrix containing 643 numeric features per assessment, with chronological age as the prediction target (Fig. 1h). Demographic variables (sex, race, education) were represented as numeric covariates in the exported matrix. For interpretability and figure generation, features were grouped into physiologically and task-grounded categories using ordered partial-string matching against feature names. Each feature was assigned to the first category whose string list matched; features with no matches were assigned to “Spectral”. Categories and match strings were: ERP Inhibit [“INHIBIT”, “CZ_Pe”, “CZ_ERN”, “NOGO-INCONGRUENT”, “NOGO-CONGRUENT”]; ERP Undistracted [“_UNDISTRACTED”, “FLANKER-CONGRUENT”]; ERP Distracted [“_DISTRACTED”, “FLANKER-INCONGRUENT”]; Gamma [“gamma”]; Delta [“delta”]; Theta [“theta”]; Alpha [“alpha”, “iaf”]; Beta [“beta”]; Behavioral [“RT-”, “AC-”, “AC_”, “RT_”]; Composite [“c_”]; Demographics [“race”, “sex”, “education”]. Definitions were: ERP Inhibit, ERP biomarkers from correct inhibit trials as well as error-related ERP biomarkers; ERP Undistracted, ERP biomarkers from correct undistracted (congruent) trials; ERP Distracted, ERP biomarkers from correct distracted (incongruent) trials; Gamma/Delta/Theta/Alpha/Beta, EEG band-annotated biomarkers; Behavioral, reaction time and accuracy measures; Composite, engineered interaction measures; Demographics, race/sex/education; Spectral, other spectral measures including broadband slope, total power, or median frequency.

### Inclusion criteria and data cleaning for model development

Model development used the cross-sectional dataset exported to a single table with one row per assessment and one column per feature. Chronological age was coerced to numeric and filtered to include adult ages 18.0–89.99 years. Rows with missing values in age or any feature column were excluded. Feature columns were restricted to numeric dtype, preserving the original column order of the export. The final ordered list of feature names was stored alongside the trained preprocessing objects and model parameters to ensure exact feature alignment during inference.

### Train-test splitting

Data was split into training and held-out test sets (80:20) using a fixed random seed (random_state = 52). To stabilize age representation across the split, an age-stratified splitting procedure was used when feasible: chronological ages were sorted and assigned to 18 bins using a fixed bin-assignment vector, and the split was stratified by these bin labels if all bins contained at least two samples. All downstream preprocessing steps were fit exclusively on the training data.

### Preprocessing pipeline

The preprocessing pipeline consisted of: (i) variance filtering (VarianceThreshold; threshold = 1e−7) fit on the training set; (ii) per-feature percentile clipping using bounds estimated from the training set (0.5th and 99.5th percentiles) and applied identically to training and test data; (iii) Yeo–Johnson power transformation with internal standardization (PowerTransformer; standardize = True) fit on the clipped training data and applied to the clipped test data; and (iv) an additional z-scoring step (StandardScaler) fit on the transformed training data and applied to the transformed test data. For longitudinal inference, the fitted variance filter, clipping bounds, transformer and scaler were serialized and reused unchanged.

### BrainYears model architecture

BrainYears prediction used a two-stage model that combines a regularized linear component with a nonlinear residual corrector. In stage 1, an ElasticNet regression model was fit to predict chronological age from the scaled feature matrix using alpha = 0.325, l1_ratio = 0.235, selection = “random”, max_iter = 100000, tol = 1e-5, and random_state = 52. In stage 2, residuals were computed on the training set as (chronological age - ElasticNet prediction). A GradientBoostingRegressor was trained to predict these residuals from a concatenated design matrix consisting of the raw clipped features (prior to power transform and standardization) together with the ElasticNet prediction as an additional column. Gradient boosting hyperparameters were: n_estimators = 185, learning_rate = 0.0175, max_depth = 2, subsample = 0.5, min_samples_leaf = 60, loss = “huber”, and random_state = 52. Final raw age predictions were computed as the sum of the ElasticNet prediction and the gradient-boosted residual prediction.

### BrainYears bias-correction procedure

To mitigate regression-to-the-mean effects, age-dependent bias was modeled and removed after the two-stage prediction. A polynomial basis (PolynomialFeatures, degree = 3, include_bias = False) was fit on training-set raw predictions, and a linear regression model was fit to predict the training-set prediction error (raw predicted age - chronological age) from these polynomial features. Corrected predictions were obtained by subtracting the predicted bias term from the raw prediction. Importantly, the bias model was a function of predicted age (not chronological age) and was trained only on the training split; the fitted polynomial transform and bias regressor were applied unchanged to the held-out test set and to all longitudinal analyses.

### BrainYears Model evaluation

Model performance was evaluated on the held-out test set using mean absolute error (MAE) and Pearson correlation between corrected predicted age and chronological age (Fig. 2a). Additional diagnostic plots included calibration curves (Fig. 2c), MAE stratified by age bin (Fig. 2d), and residual distributions across age (Fig. 2e). All reported evaluation metrics and diagnostics used predictions generated with a single frozen train-test split (random_state = 52) and preprocessing fit on training data only.

### Domain-level interpretability analyses

To summarize model structure at the domain level (Fig. 3a–h), analyses were performed using the feature-category mapping described above. Category-level effects included: (i) distributions of feature-age associations computed per feature and summarized by category (Fig. 3a); (ii) category permutation importance computed by shuffling all features within a category together and quantifying the increase in prediction error (Fig. 3b); (iii) ElasticNet coefficient directionality summarized as the fraction of positive versus negative coefficients per category (Fig. 3c); and (iv) band-focused summaries of age-associated effects for canonical EEG frequency bands (delta, theta, alpha, beta, gamma) (Fig. 3d). Where stability metrics were shown (Fig. 3f), selection stability was defined as the frequency with which features were retained after the variance filter across resampled model fits under identical preprocessing and hyperparameter settings.

### Longitudinal cohorts and intervention adherence

Longitudinal application of the clock was performed on two cohorts using paired assessments per participant. Participants were indexed by missionId, and assessmentNumber provided the within-person ordering of assessments. For each assessment, chronological age and the full feature set required by the trained bundle were required; rows with missing age, missionId, assessmentNumber, or any required feature columns were excluded.

#### Intervention cohort

Participants were selected if their first interaction with Sens.ai was completion of the Sharp Mind Mission protocol, they were at least 40 years old (participants 41 - 82 years old, 55 Average), and they met an adherence criterion of at least 4.5 sessions per week (7 sessions per week on average). Participants completed their first assessment within 6 days pre-intervention and a second assessment after completion within 9 days post-intervention. The interval between assessments ranged from 33 to 92 days (average 56 days). The intervention cohort consisted of 28 participants (3 female, 25 male). Participants completed a minimum of 20 sensorimotor rhythm neurofeedback sessions and 20 Peak Alpha with Gamma neurofeedback sessions at an average pace of 4.5 or more sessions per week; participants were allowed to complete additional neurofeedback sessions and were encouraged to complete heart rate variability biofeedback and transcranial photobiomodulation booster sessions. On average, participants completed 21 sensorimotor rhythm neurofeedback sessions, 20 Peak Alpha with Gamma neurofeedback sessions, 12 heart rate variability biofeedback sessions, and 18 transcranial photobiomodulation sessions between assessments.

#### Control cohort

Participants were included if they initiated a protocol but completed no more than 5 headset sessions (3 sessions on average). Ages ranged from 25 to 65 years (average 45 years). Assessments were 11 to 84 days apart. This cohort consisted of 22 participants (7 female, 15 male).

### Longitudinal outcome definition and statistics

For each assessment, brain-age delta was defined as bias-corrected predicted brain age minus chronological age. Within each participant (missionId), pre/post change in brain-age delta was computed as the difference between the final and initial assessments (ordered by assessmentNumber): Δdelta = delta_last − delta_first. Negative values correspond to improvement (a shift toward a younger predicted brain age relative to chronological age). Group summaries included the mean Δdelta and the percentage of participants showing improvement (Δdelta < 0) (Fig. 2f–h).

### Software and reproducibility

All analyses were implemented in Python using NumPy, pandas, SciPy, and scikit-learn. Model training outputs were serialized as a single joblib “bundle” containing the ordered feature list, fitted preprocessing objects, trained ElasticNet and gradient boosting models, and fitted bias-correction components. Longitudinal inference reused this frozen bundle to ensure identical preprocessing and prediction logic across cohorts. Code will be made available upon request.

## Acknowledgements

This research was funded by Hevolution Foundation to the Buck Institute for Research on Aging (HF-PART-23-1422047).

## Author contributions

C.J., P.T., and E.V. conceived the study. C.J. preprocessed the data and provided the curated dataset. S.L. and C.J. developed the model and performed the computational analyses. M.S.-K. and E.V. contributed to conceptual framing and interpretation of the results. S.L. wrote the manuscript and generated the figures. All authors contributed to manuscript revision and approved the final version.

---

**Supplementary Figure 1.**
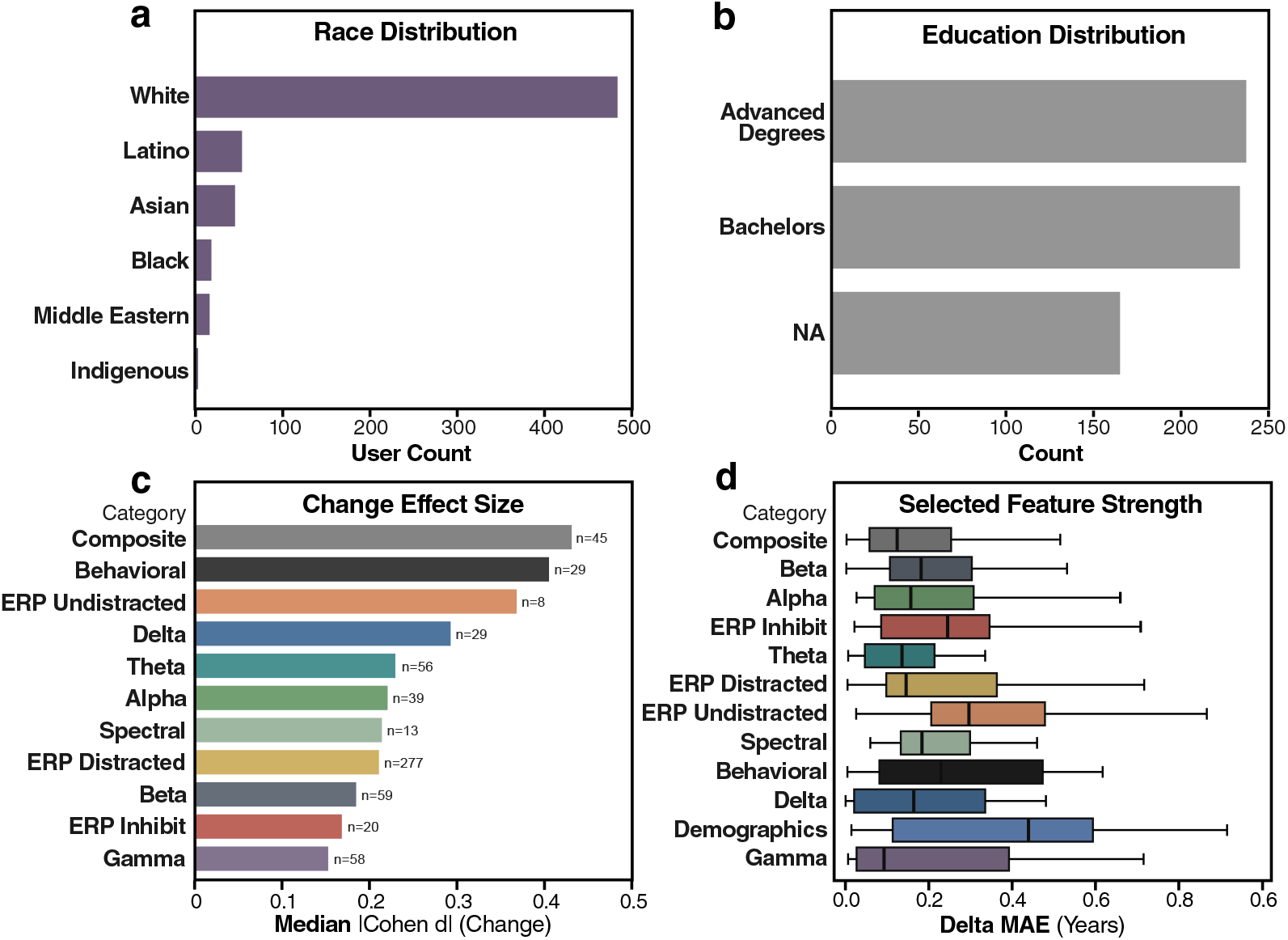
Cohort demographics and domain-level change metrics. **a**, Self-reported race distribution. **b**, Highest educational attainment distribution. **c**, Median within-participant Cohen’s d for pre–post change stratified by feature category. **d**, Distribution of absolute model coefficient magnitudes by feature category.

