## Supplemental Figures for "BrainYears: A functional EEG-based brain age clock enables intervention-ready measurements of brain aging"

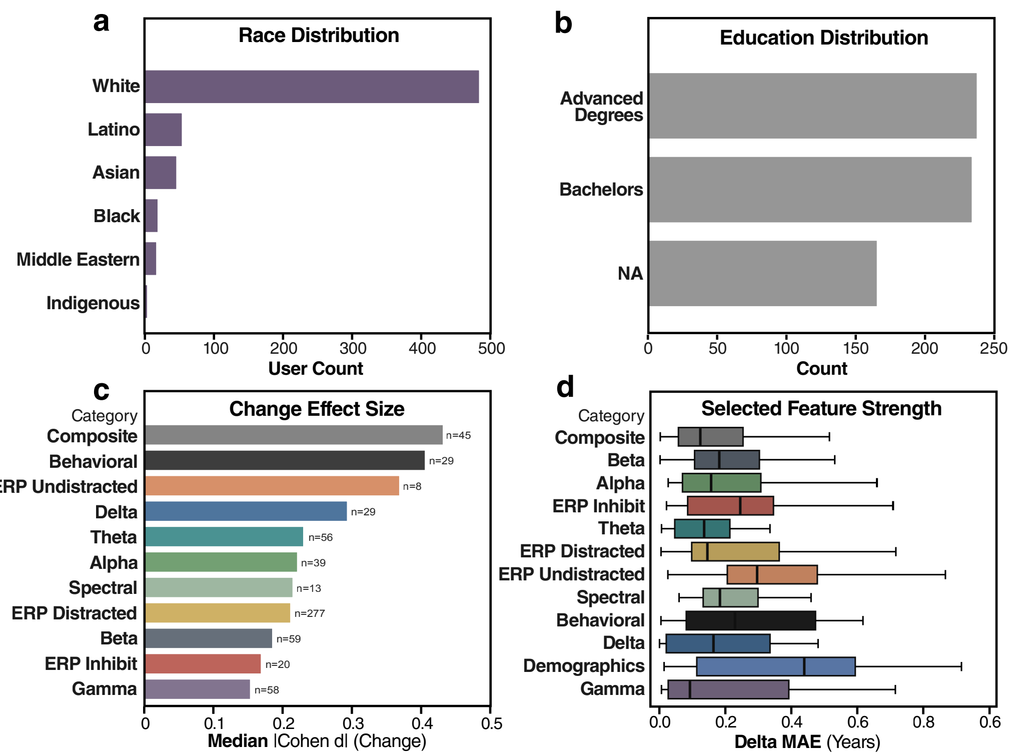


**Supplementary Figure 1 | Cohort demographics and domain-level change metrics.**
**a,** Self-reported race distribution. **b,** Highest educational attainment distribution. **c,** Median within-participant Cohen's d for pre–post change stratified by feature category. **d,** Distribution of absolute model coefficient magnitudes by feature category.
